# Subdomains of Endophilin-NBAR Can Synergistically Drive Membrane Remodeling and Facilitate Controlled Membrane Scission

**DOI:** 10.64898/2026.02.19.706845

**Authors:** Jeriann R Beiter, Feng-Ching Tsai, Patricia Bassereau, Gregory A. Voth

## Abstract

The NBAR-domain containing protein endophilin, as a major player in many endocytic pathways, has offered considerable insight into BAR-domain driven membrane remodeling. However, understanding the interaction of the different subdomains of endophilin and their abilities to sense and generate negative Gaussian curvature are yet unanswered questions, with significant implications for the mechanisms and regulation of unconventional endocytic pathways. Using coarse-grained molecular dynamics simulation, we demonstrate the synergistic remodeling capabilities of the NBAR remodeling unit, as well as its ability to sort to and generate membrane regions with negative Gaussian curvature. We find that the assembly of NBAR scaffolds at regions of negative Gaussian curvature facilitate membrane hemifission in dynamic bud formation. These insights provide an additional role for endophilin scaffolds in endocytosis, as well as emphasizing the importance of developing new ways to study negative Gaussian curvature.

**STATEMENT OF SIGNIFICANCE:** This work provides deeper insight into the composite membrane remodeling abilities of NBAR domains in peripheral membrane proteins and their sorting to negative Gaussian curvature. Theis work also explicitly models at the molecular level the Ω-shaped membrane geometry, connecting endophilin mechanics to its physiological function in endocytosis.

## INTRODUCTION

Endocytosis is a class of cellular processes critical for homeostasis, where a portion of the plasma membrane is curved and eventually severed to create a separate, internalized vesicle, which enables, e.g., cargo and nutrient uptake, receptor recycling, and plasma membrane tension regulation. A variety of endocytic pathways exist, primarily defined by the cargo type (IL2Rβ uptake), the major membrane remodeling proteins involved (caveolin-mediated, clathrin-mediated (CME)), or even the relative speed of the process (ultra fast (UFE) and fast endophilin mediated (FEME))(1-3). However, one commonality that unites these processes, and indeed most plasma membrane remodeling, is their reliance on Bin/Amphiphysin/Rvs (BAR) domain containing peripheral membrane proteins.

BAR domains consist of a three-helix coiled-coil bundle that dimerize via a hydrophobic core and have positively charged residues at the ends of bundle to facilitate binding to negatively charged lipids. Of the BAR domain proteins involved in endocytosis, endophilin is perhaps the most ubiquitous. First characterized in the context of synaptic vesicle endocytosis(4,5), endophilin has since become a canonical example of an NBAR-domain protein, where the N designates an N-terminal amphipathic helix directly connected to the BAR domain; in the case of endophilin-A1, this N-terminal amphipathic helix is referred to as Helix 0, or H0 for short. In addition to the NBAR domain, endophilin-A1 also has a C-terminal SH3 domain that recruits other downstream effector proteins such as dynamin(6-8), as well as a 41 residue disordered linker domain connecting the BAR and SH3 domains.

Previous work has identified endophilin as a potent membrane remodeling agent, both *in vivo* and *in vitro*. In particular, endophilin has been shown to generate membrane tubes from the plasma membrane and giant unilamellar vesicles (GUVs), as well as reshape small unilamellar vesicles (SUVs) into tubes with well-defined diameters of ∼25 nm(9,10). Even the H0 amphipathic helix alone is capable of remodeling supported lipid bilayers (SLBs) into SUVs(11). In addition to its remodeling capacities, endophilin is also sensitive to preformed membrane curvatures, on the order of tens of nanometers(12,13).

These observations have led to questions of how endophilin and other BAR domain proteins are able to generate and be recruited to (or “sense) nanometric membrane curvatures, with various claims centering on different subdomains of endophilin and their hypothesized remodeling mechanisms. The hypothesized molecular membrane remodeling mechanisms are insertion, scaffolding, and crowding(14-16), which in the case of endophilin have each been mapped to the amphipathic helix H0, the BAR domain, and the unstructured linker and the SH3 domain, respectively. While some comparative analysis of the endophilin subdomains in remodeling has been conducted and emphasized the role of the H0 insertion in generation and sensing of membrane tubes, these studies have primarily focused on conditions not representative of physiological conditions such as high protein concentration regimes or membranes composed almost entirely of negatively charged lipids such as DOPS(10,17,18). Additionally, the importance of the unstructured linker and SH3 domains in driving membrane curvature generation have only been studied in the context of similar high protein concentration regimes (19-21).

Furthermore, all of the previous membrane geometries studied either experimentally or computationally have had zero (SLBs, flat sheets) or positive (SUV/GUVs, pulled tubes, membrane tube and vesicle simulations) Gaussian curvature. This is attributable to the difficulty of generating bilayer surfaces with negative Gaussian curvature, particularly at the nanometric scale. However, imaging and continuum modeling of endocytic events have consistently shown an Ω-shaped membrane geometry as a key intermediate between the initially flat surface and scission, where the neck has a catenoid geometry(22-25). Understanding how endophilin and indeed other peripheral membrane proteins sense and generate negative Gaussian curvature on membranes is a key challenge that has remained largely unaddressed.

In this work, we have probed these questions using a combination of coarse-grained (CG) molecular dynamics (MD) simulation to understand the interaction of endophilin with model membranes. By simulating piecewise various endophilin constructs on both flat and tube membranes, we systematically explore the synergistic membrane-remodeling capabilities of the H0 and BAR domains as well as how membrane-mediated effects drive their collective assembly. We further explore the ramifications of this synergy by modeling undulating membrane tubes with negative Gaussian curvature to explore whether the endo-NBAR domain is recruited to and similarly assembles on these membranes. Finally, we simulate the Ω-shaped geometry similar to those observed in endocytic buds to compare the differential recruitment and dynamic impact of endophilin networks on membrane scission.

## METHODS

### Atomistic Molecular Dynamics

The initial structure of the BAR domain of endophilin-A1 was taken from PDB ID 1ZWW(26), with the missing N-terminal amphipathic helices added with the structure editor feature in ChimeraX(27). The resultant structure was initially positioned 1 nm above a 45×45nm^2^ symmetric membrane with composition of 80:20 DOPC:DOPS and solvated with a 0.15M KCl solution, prepared with CHARMM-GUI(28,29). After a 1.87 ns equilibration in the constant NVT ensemble, production simulations were run in triplicate for 450 ns total (150 ns each) with a 2.0 fs timestep in the constant NPT ensemble using the CHARMM36m forcefield in GROMACS2020(30,31). Previous examples of atomistic MD simulations of N-BAR domain proteins interacting with membranes can be found in refs. (32-34)

The initial structure of the SH3 domain and unstructured linker were modeled with AlphaFold2(35). The resultant structure was solvated with a 0.15M KCl solution in a 10.7×10.7×10.7 nm^3^ box, and equilibrated for 0.75 ns before running 100 ns for production.

### Coarse-Grained Molecular Dynamics

The production portion of the atomistic MD simulations were mapped into a resolution with one CG site or “bead” representing one amino acid and placed at the α-carbon position. A heteroelastic network model (hENM)(36) was developed using OpenMSCG to generate the intramolecular protein potentials by matching the atomistic fluctuations from the mapped trajectory(37). The lipids were modeled using a four CG site linear representation corresponding to a head, interfacial, and two tail CG sites, with the intramolecular bond and angle potentials represented harmonically with kappa values of 15.4 kcal/mol/Å and 1.23 kcal/mol/degree and r_0_ values of 7.5 Å and 180^0^, respectively(38). Intermolecular lipid-lipid potentials were modeled with a piecewise cosine potential(39), where B is instead set as 0.18 for DOPC and 0.19 for DOPS, and r_0_ set as 8.0 and 7.5 Å respectively. The protein-lipid intermolecular potentials were generated by fitting the same potential form to numerical potentials first generated via force matching using OpenMSCG, which allowed for the generation of smoother potentials with consistent forms. Only the head and interfacial sites of the CG lipids interact with the protein sites, consistent with the sampling in our atomistic simulations.

Coarse-grained simulations were conducted with the LAMMPS MD software(40) with a 25 fs timestep in the NPT ensemble, with coupling in the xy plane. All CG simulations had a membrane composition of 80:20 DOPC:DOPS. All simulations are initialized with proteins randomly distributed 0.5 nm away from membrane surface. A summary of all CG simulations can be found in Table S1 of the Supporting Information. For the bud generation simulations, a resistive tension of -0.15 atm was applied to prevent excessive deformation at the boundaries due to box size changes. Analysis and visualization conducted using the Python package MDAnalysis and Visual Molecular Dynamics (VMD)(41,42).

## RESULTS AND DISCUSSION

### Endo-NBAR as a Composite Unit is Most Effective at Binding and Remodeling Membranes

To explore how the different subdomains of endophilin contribute to its membrane remodeling capacity at the collective scale, we simulated several different subconstructs of our endophilin CG model at a molecular level (Figure 1A). While previous MD simulations have explored the impact of an individual endophilin dimer, or even a small handful of them, none have explicitly modeled comparatively how each subdomain of endophilin drives large-scale membrane remodeling(12,18,32,43-45). In particular, we wanted to explore whether there was a synergistic relationship between the H0 helix, the BAR domain, and the unstructured SH3 linker region and their respective proposed remodeling mechanisms of insertion, scaffolding, and crowding.

**Figure 1:**
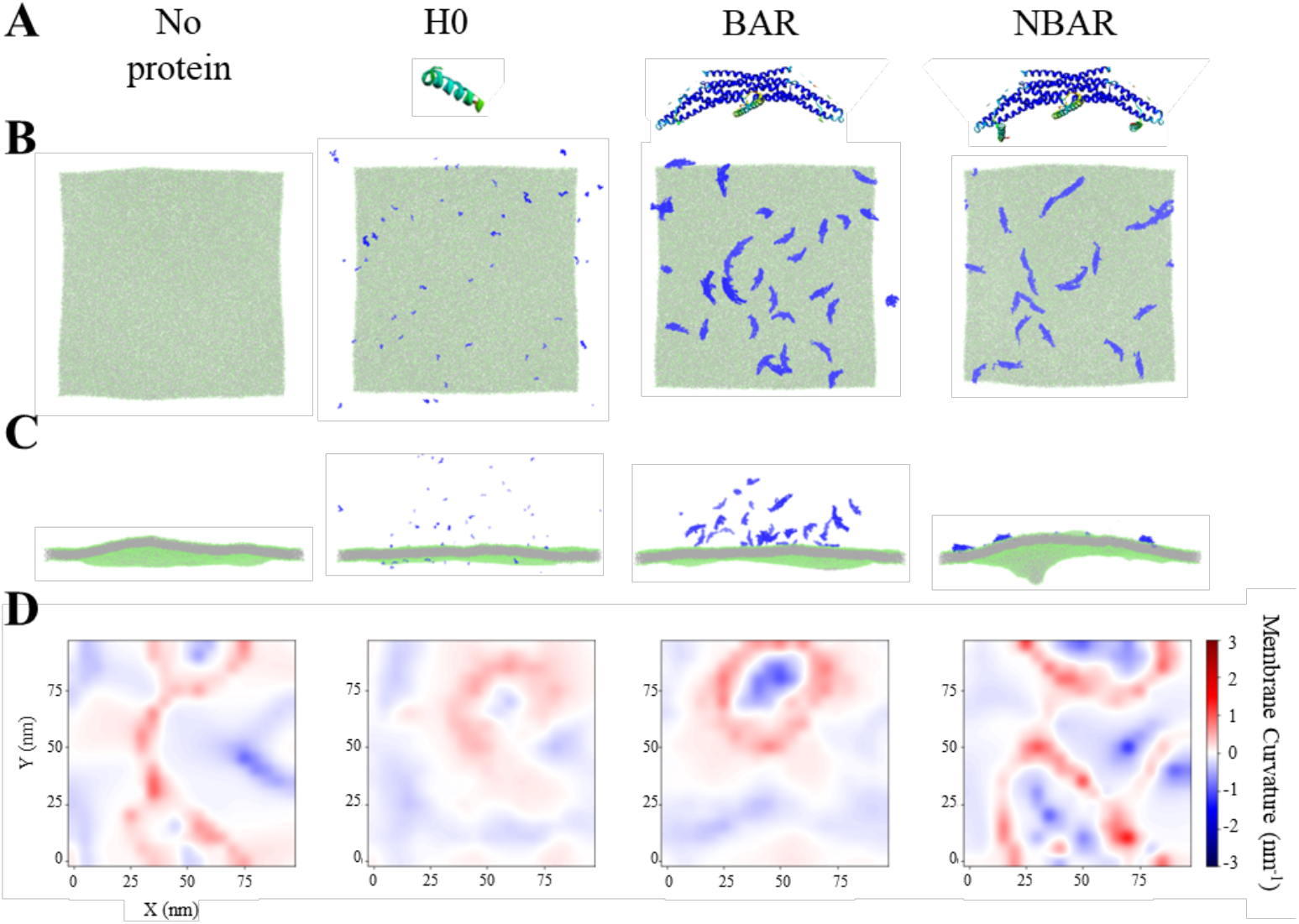
The H0 and BAR domains of endophilin have synergistic membrane curvature generation effects on initially flat membranes. **(A)** Structural representation of the different constructs simulated (left to right): no protein (undecorated), H0 helix, BAR domain, and composite NBAR domain. Structures visualized with ChimeraX(27). **(B)** Top and **(C)** Side view snapshots of CG systems after 100 10^6^ CG timesteps (CGts) from random initial protein placement on 100×100nm^2^ membranes at 10% surface coverage. Various protein constructs are represented in blue in the same order as (A), with the lipid head sites shown in green and tail sites shown in silver, visualized with VMD(42). **(D)** Mean membrane curvature of upper (protein-bound) leaflet after 100 10^6^ CGts, represented from left to right in same order as (A). Membrane curvature calculated with MDAnalysis package MembraneCurvature(41).

Previous work has demonstrated that the NBAR domain of endophilin is quite potent on its own as a remodeling unit, and that a relatively low surface coverage (up to 35%) is necessary to effect changes in membrane shape(12). With this in mind, we initially investigated the effect of the NBAR dimer remodeling on 100nm x 100nm flat sheets at surface coverage of 10% (32 dimers); for comparison, we kept this consistent with the other constructs as well with 32 BAR dimers, 32 full length endophilin (FL-endo) dimers, and 64 individual H0 helices (Figures 1,S1; Table S1). While all protein constructs are initially randomly placed above the membrane surface, only the NBAR and FL-endo constructs consistently and stably bind to the membrane over the course of the simulation. The BAR and H0 helices can and do associate with the membrane surface, but the majority reside in the “bulk” space unassociated with the membrane. As a result, the effective curvature generation of the BAR and H0 constructs is minimal and similar in magnitude to the control membrane without any proteins (Figure 1D). By contrast, the NBAR and FL-endo produce significant changes in the membrane curvature, and they are even ordered into string-like assemblies by the membrane, consistent with previous observations(46,47). The synergistic impact of the H0 and BAR domain bound together as a compact NBAR unit as a more dramatic effector of membrane remodeling has been previously reported, with much higher concentrations of BAR domain or H0 helix alone necessary to demonstrate similar levels of remodeling to the NBAR construct(12,48). By contrast, the addition of the disordered linker and SH3 domain to the NBAR domain in the FL-endo does not appear to drive a significantly greater remodeling mechanism induced by the flexibility of the linker (Figure S1). Crowding has been identified as important in the membrane remodeling capabilities of amphiphysin, another BAR-domain containing protein, but the unstructured linker region of amphiphysin is longer, 382 amino acids in length, compared to that of endophilin (41 amino acids)(20). We therefore note, in agreement with previous literature, endophilin alone at this surface density is insufficient to generate liquid-liquid phase separation mediated membrane remodeling due to the length of its linker region (19-21).

To better understand how these results may translate to the tubular geometries that endophilin is most often associated with, we translated our construct study onto 60nm periodically repeating tubes with diameters of 30nm (Table S1, Figure 2A). This diameter is slightly larger than the typical curvature generated by endophilin of 25 nm(10), allowing us to observe whether various constructs not only bind but also induce a curvature closer to their preferred geometry. Similar to the effects observed on a flat membrane, both the H0 and BAR constructs do not appreciably remodel the membrane tube, again due to reduced binding efficacy. By contrast, the NBAR domain binds to and remodels the tubular membrane surface, inducing the most dramatic sustained changes in the major and minor tube radii compared to the control membrane or systems with the H0 or BAR domain constructs (Figure 2B). While there is a dynamic range in the tube anisotropies for each system, the ellipticity, defined as the ratio of the major and minor radii, of the membrane with NBAR constructs is on average higher than the other systems (Figure S2). The higher average ellipticity observed in the H0 and BAR domain constructs compared to the control trajectory are also likely due to their transient binding. Interestingly, the dip in ellipticity of the NBAR-containing system at z = 25nm is due to an overall constriction of the membrane, as quantified in Figure S2.

**Figure 2:**
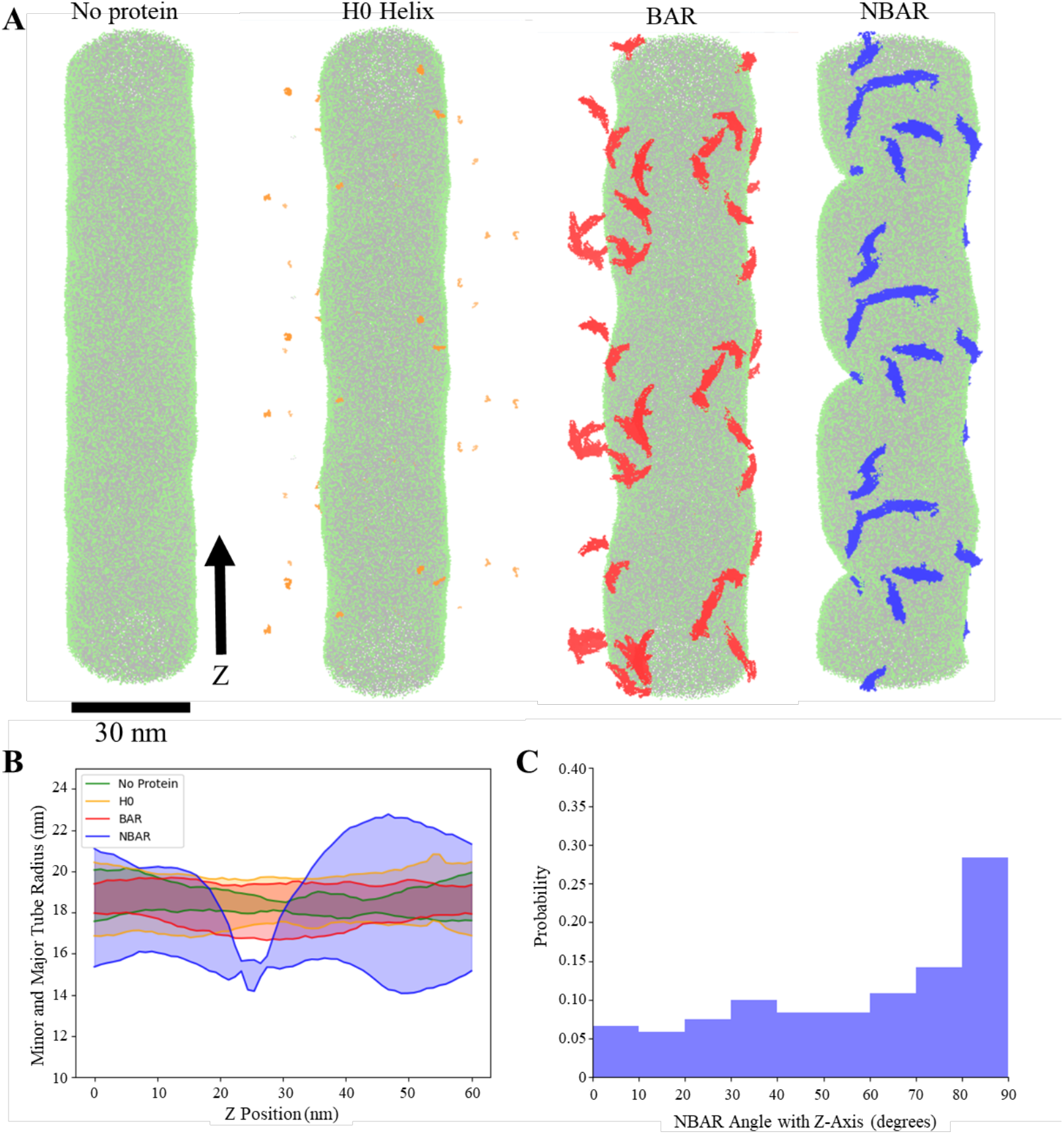
The NBAR domain is effective at remodeling tubular membranes and is itself subsequently reorganized. **(A)** Side view snapshots of CG systems with 30 nm inner diameter membrane tubes after 50 10^6^ CG timesteps (CGts) from random initial protein placement. The H0 helix sites are represented in orange, the BAR sites are represented in red, and the NBAR sites are represented in blue, with the lipid head sites shown in green and tail sites shown in silver for all systems. Visualized in VMD with ±Z periodic images to show curvature. **(B)** Minor and major tube radii after 50 10^6^ CGts. Radii calculated based on CG tail bead positions subdivided into 1 nm wide slices for membranes with no protein (green), with 24 NBAR dimers (corresponding to 10% surface coverage; blue), with 24 BAR dimers (red), and with 48 H0 helices (orange). **(C)** Histogram of orientation of NBAR protein domain with respect to central z-axis, calculated from 45-50 10^6^ CGts. 0 degrees represents the major axis of the NBAR domain aligning parallel with the major axis of the tube (z-axis), and 90 degrees represents perpendicular alignment.

The membrane constriction corresponds spatially to membrane-mediated aggregation of the NBAR constructs into linearized assemblies of 3-5 dimers (Figure 2A, right). Similar to experimentally resolved structures of endophilin on tubular membranes, we see that the preferred alignment of NBAR dimers is perpendicular to the tube axis at this membrane diameter (Figure 2C)(10). As we observe this effect at significantly reduced surface coverage compared to the experimental structures, we affirm that the endophilin NBAR curvature preference is based on membrane geometry and not protein-protein interactions.

### Endo-NBAR is Recruited to and Generates Negative Gaussian Membrane Curvature

While the effect of NBAR mediated remodeling has previously been studied – primarily in the context of membrane straight tubes and flexible sheets – the effects of Gaussian membrane curvature on the recruitment and organization of endophilin remain unknown. In part, this is due to the difficulty of experimental reconstitution of stable saddle-shaped membrane curvatures, particularly at the dimensions observed during endocytosis(23,25).

To explicitly probe how endophilin NBAR organize on systems with negative Gaussian curvature, we set up a membrane tube with an explicitly formed catenoid shape, with internal harmonic regions to maintain the membrane shape. After 50×10^6^ CG timesteps (CGts), we see that the membrane maintains the catenoid tube shape (Figure 3A), though there is some relaxation in the region with negative mean curvature in the system without NBAR (Figure S3A). By contrast, in the system with 10% NBAR initially placed near the membrane we see that the membrane curvatures are amplified, with a comparatively narrower minor radius in the region with negative mean curvature, and an overall increase in tube ellipticity (Figures. 3B, S3B). It must be noted here that “time” (and CGts) in a CG simulation is not the same as real time, as the sampling in the CG model is (as desired to achieve enhanced sampling) greatly accelerated over, for example, all-atom MD timescale(49). To associate the CG timescale with a real timescale requires that one scales the former based on a known experimental timescale for the latter real processes.

**Figure 3:**
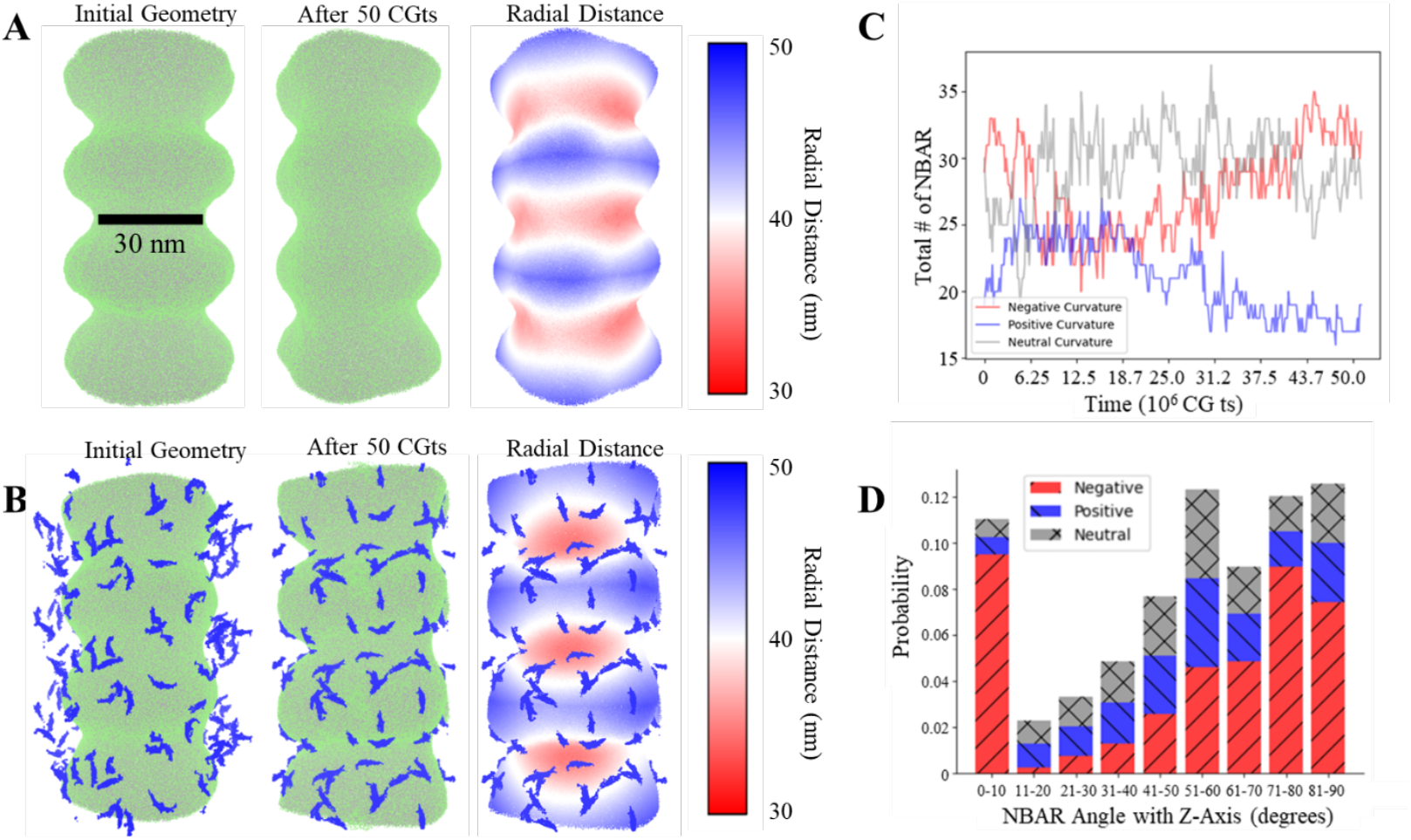
The NBAR domain is recruited to negative Gaussian curvatures and creates local lipid reservoirs. Side view snapshots of Gaussian curvature tubes without **(A)** and with 10% NBAR **(B)** designating initial position (left) and after 50×10^6^ CGts (center). The NBAR sites are represented in blue, with the lipid head sites shown in green and tail sites shown in silver for both systems in left and center; lipids are colored by radial distance from the Z-axis in the right panels (red – smaller radial distance; blue – larger radial distance). Visualized in VMD with ±Z periodic images to show curvature. **(C)** Timeseries of NBAR sorting to areas of negative (red), positive (blue), and neutral (gray) mean curvature. **(D)** Histogram of orientation angles with Z-axis of NBAR dimer, taken over last 5 10^6^ CGts of simulation. Bars are colored and textured based on their mean curvature sorting, as in (C).

By defining spatially equally sized regions of initially positive, negative, and neutral mean curvature, we can track over the course of the simulation to where NBAR domains are recruited. While the NBAR domain proteins are evenly distributed in space initially and take about 6×10^6^ CGts to effectively bind to the membrane, we see a redistribution and subsequent plateau of the distribution that heavily favors the regions of negative and neutral mean curvature over positive mean curvature (Figure 3C). When considering the orientation of the NBAR dimers relative to the Z-axis, we see a general trend towards an antiparallel alignment (closer to 90 degrees) for all sorting, consistent with Figure 2C. However, there is a noted exception at 0 degrees, indicating a significant proportion of NBAR dimers in the negatively curved region are almost exactly parallel to the Z-axis (Figure 3D). Given that this behavior is not observed in the straight tube (Figure 2A), whose diameter corresponds most closely to the region of negative mean curvature, we conclude this is an effect of the catenoid geometry.

As calculating the Gaussian curvature of an irregular 3D surface is nontrivial, we have done so here by partitioning the membrane into two Monge surfaces split at the YZ (Figure 4A) or XZ planes. We then calculate the curvature for each surface separately (Figure 4B-C, S4). At greater radial distances in x or y (for the YZ plane and XZ plane surfaces, respectively), there is an underestimation of the curvature due to the binning procedure averaging over a greater number of lipids as we approach the edge of the surface, which is why both the YZ and XZ surfaces have been calculated, to best approximate the overall surface Gaussian curvature. Even with this in mind, we note a dramatic increase in the local Gaussian curvature generation with 10% NBAR dimers, both negative and positive. This increase is particularly apparent in the regions of globally high Gaussian curvature, where the catenoid tube radius is at a minimum or maximum. Further, the fact that the local Gaussian curvature at 10% surface coverage of NBAR dimers has a more “patchwork” spatial representation indicates significant local reorganization of the underlying lipids.

**Figure 4:**
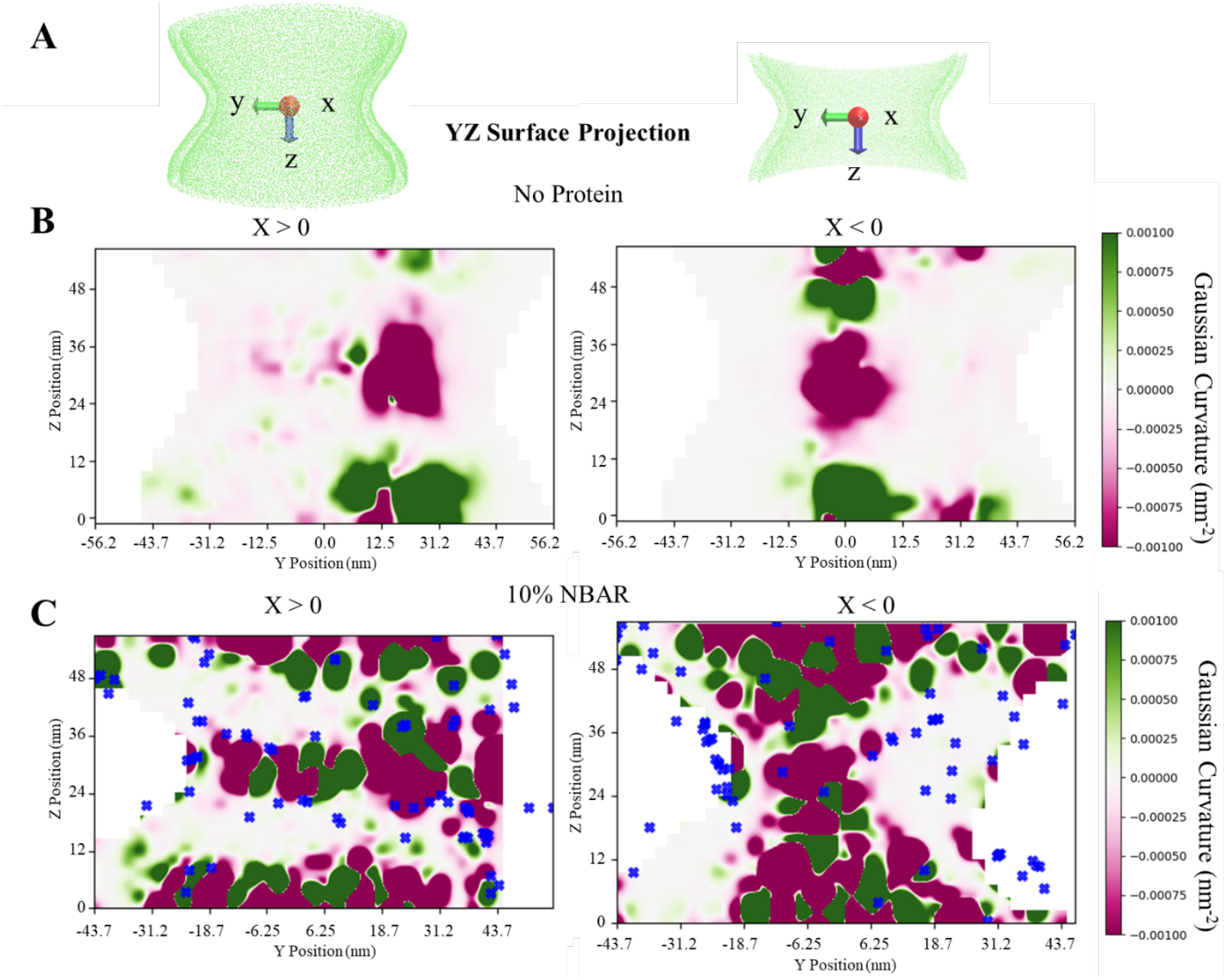
Endo-NBAR generates substantial local Gaussian curvature on catenoid tubes. **(A)** Representation of catenoid tube divided by the YZ plane for X > 0 (*left*) and X < 0 (*right*). **(B)** Front (X > 0, *left*) and back (X < 0, *right*) of catenoid tube membrane without NBAR protein projected onto yz plane after 50 10^6^ CGts, with Gaussian curvature calculated over 2.5×2.5nm^2^ bins. **(C)** Front (X > 0, *left*) and back (X < 0, *right*) of catenoid tube membrane with 78 NBAR dimers (corresponding to 10% surface coverage) projected onto yz plane after 50 10^6^ CGts, with Gaussian curvature calculated over 2.5×2.5nm^2^ bins. NBAR dimer tip positions shown as blue x’s. Gaussian curvature calculated with MDAnalysis package MembraneCurvature.

One consequence of this generation of membrane curvature is the reorganization of lipids near the NBAR dimers. This reorganization is clear when comparing the lipid-lipid head group radial distribution functions (RDF) for the catenoid tube without NBAR dimers, with NBAR dimers, and the lipids that are within 1nm of an NBAR dimer (Figure S5). We note that when comparing the lipid organization via the RDF with and without NBAR dimers overall, we see a decrease in the RDF peak magnitude from 32 to 17, reflecting a moderate decrease in the overall lipid packing for the membrane with NBAR dimers. This is likely attributable to the global increase in packing defects due to the significant local curvature changes. However, when looking only at lipids in the area local (within 1 nm) of an NBAR dimer, we see a large increase in the RDF peak magnitude to 91, indicating a significant increase in tight lipid head group packing compared to the overall membrane. Taken with the increase in regions of negative Gaussian curvature, we find that NBAR dimers can create local lipid “reservoirs” that are tightly packed around the dimer.

### Endo-NBAR Dynamically Assembles at the Neck of an Endocytic-Mimicking Geometry

While we have so far offered insight into NBAR-mediated membrane remodeling of relatively static membrane geometries, we aspire here to further link the two by looking at dynamically driven shape changes, particularly in the omega-shaped membrane geometry relevant to endocytosis. By driving a spherical “cargo” region with a 30nm diameter through an initially flat geometry at a constant velocity (0.5 nm/10^6^ CGts), we can mimic the membrane geometries relevant to endocytosis, up to membrane scission. The active pulling of the cargo through the membrane is analogous to the role of actomyosin in endocytosis to drive the process forward, enabling the creation of an Ω-shaped geometry(50,51).

As the cargo is driven through the membrane, we note that an endocytic-bud like geometry is formed (Figure 5A, 5B). This cargo is well wrapped by the membrane, and generates a defined bud and neck. From Figure 5A, we see that there are few NBAR domains localized to the bud itself, as they have instead been largely redistributed to the neck, while the majority remain on the flat bilayer base. Linear string-like assemblies form on the flat bilayer base (red- and orange-colored dimers), similar to those observed in Figure 1B. In addition, ring-like scaffold structures form around the neck, near the base of the bud (yellow- and green-colored dimers). These ring-like structures are reminiscent of those observed by Hohendahl et al(52), but at a significantly lower surface coverage, demonstrating the potent ability of membrane curvature to act as an organizing cue for protein assemblies. This result provides further evidence that endophilin is sorted to areas of negative Gaussian membrane curvature, both statically such as in the catenoid tube or dynamically as is the case here.

**Figure 5:**
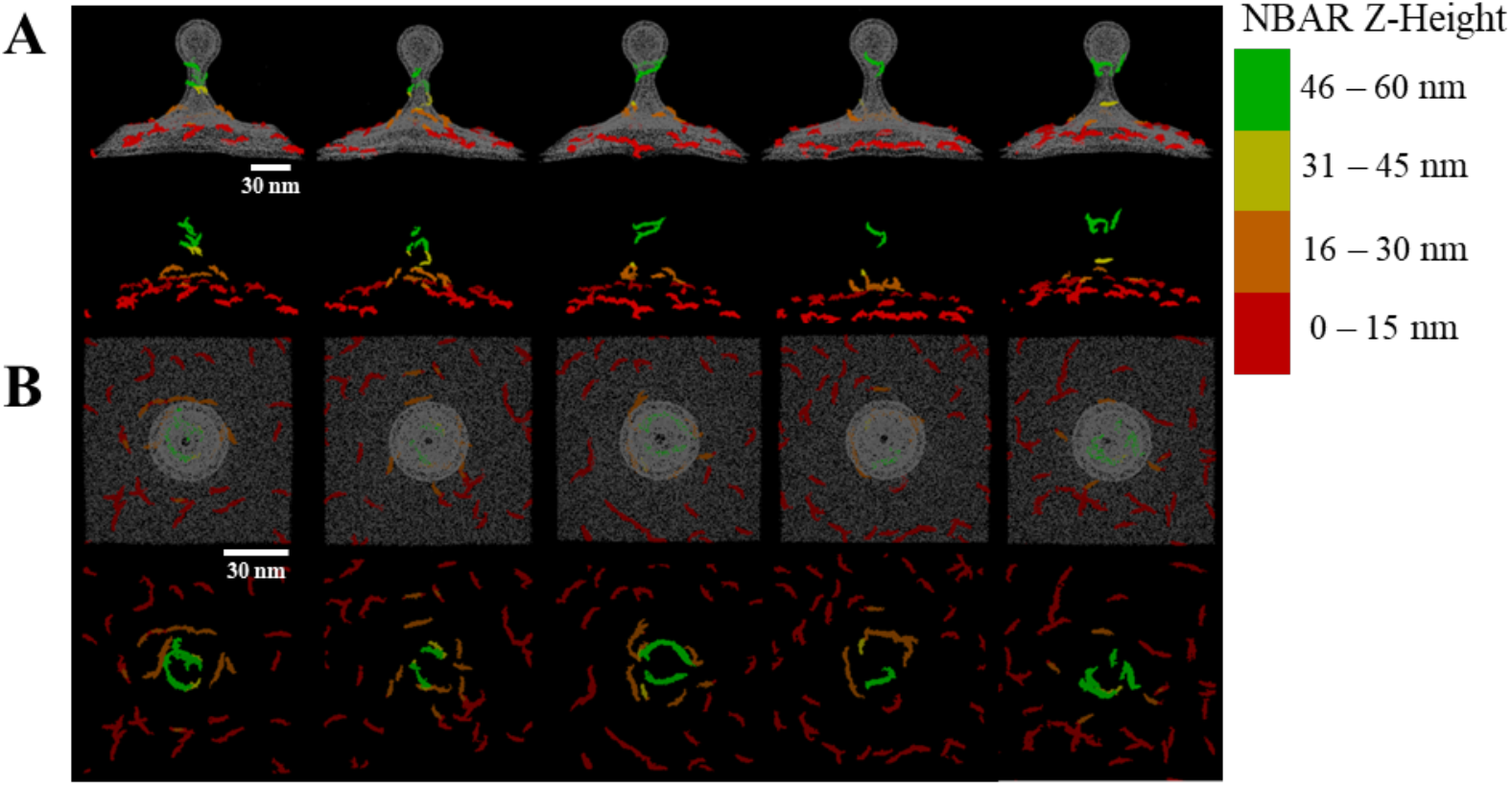
Endophilin organization at the neck of Ω-shaped geometries. **(A)** Side view of “bud-formation” simulations after 175×10^6^ CGts for all five replicates, with (*upper panels*) and without (*lower panels*) membrane explicitly shown in gray. Endo-NBAR at 10% surface coverage, colored by z coordinate. **(B)** Top view of “bud-formation” simulations after 175×10^6^ CGts for all five replicates, with (*upper panels*) and without (*lower panels*) membrane head beads explicitly shown in gray. Endo-NBAR at 10% surface coverage, colored by z coordinate. (A) and (B) visualized in VMD.

To build upon this point, we expanded further on our bud generation simulations with a system without any NBAR dimers (0% surface coverage, Figure 6A), 10% NBAR dimer surface coverage (Figure 6B), and 25% NBAR dimer surface coverage (Figure 6C). While up to this point, all simulations were done based on a 10% surface coverage of NBAR dimers, we were interested in observing how a higher concentration of NBAR dimers may influence sorting and remodeling at the membrane neck – an idea inspired in part by work suggesting that preassembled patches facilitate rapid clathrin-independent endocytic pathways(8). By continuing to drive the ‘cargo’ further in the +Z-direction, the membrane eventually narrows to the point of hemifission, detailed further in Figure S6.

**Figure 6:**
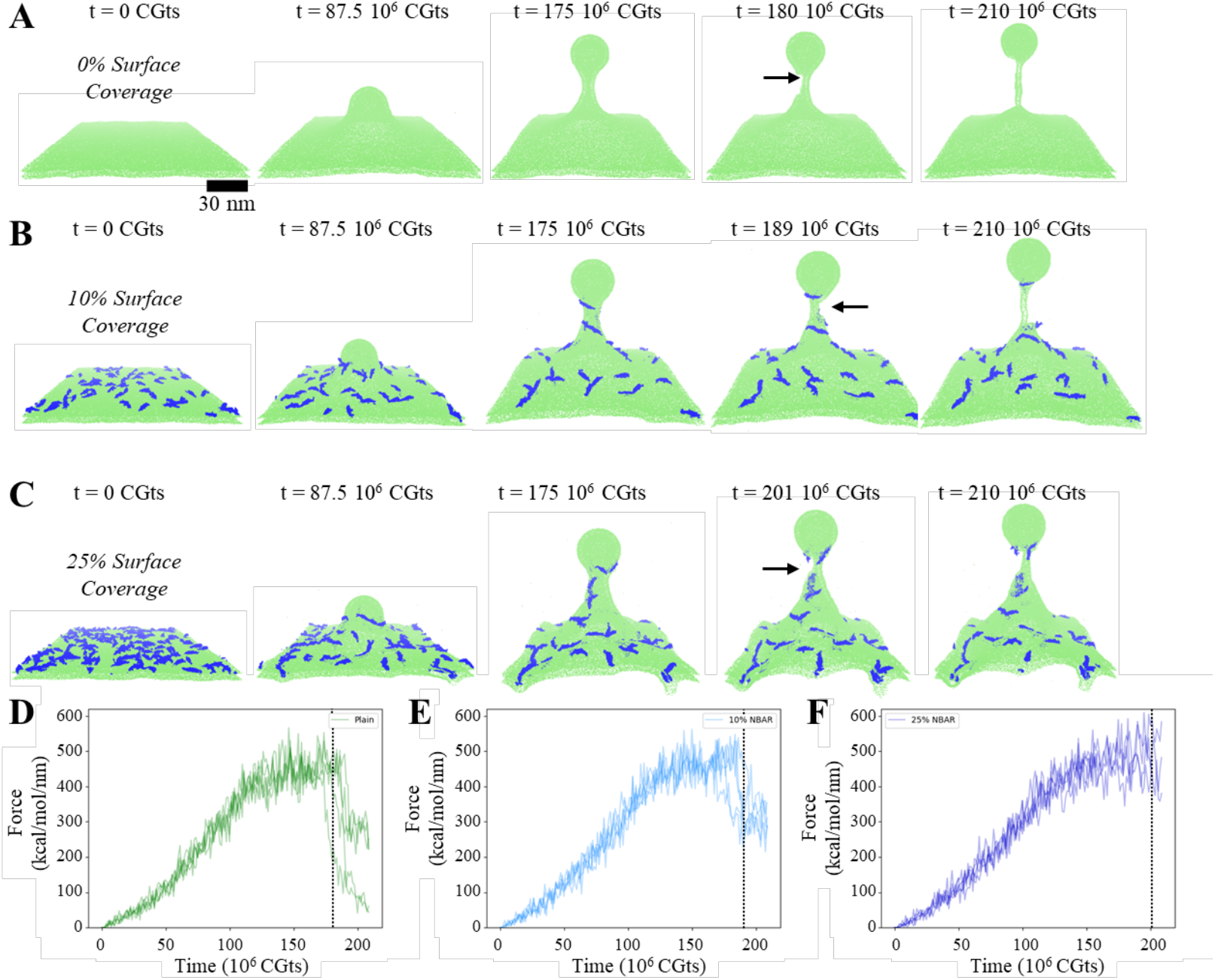
NBAR networks delay scission and prevent leakage during active transport. **(A)** Snapshots over the course of a representative “bud-formation” trajectory without NBAR at t = 0, 87.5, 175, 180, and 210×10^6^ CGts. Arrow in panel for t = 180×10^6^ CGts denotes point of membrane breakage, which is first visible in this frame. Lipid head CG sites represented in green; tails not visualized for clarity. **(B)** Snapshots over the course of a representative “bud-formation” trajectory with 10% surface coverage of NBAR at t = 0, 87.5, 175, 189, and 210×10^6^ CGts. Arrow in panel for t = 189×10^6^ CGts denotes point of membrane breakage, which is first visible in this frame. Lipid head CG sites represented in green, and NBAR CG sites represented in blue. **(C)** Snapshots over the course of a representative “bud-formation” trajectory with 25% surface coverage of NBAR at t = 0, 87.5, 175, 201, and 210×10^6^ CGts. Arrow in panel for t = 201×10^6^ CGts denotes point of membrane breakage, which is first visible in this frame. Lipid head CG sites represented in green, and NBAR CG sites represented in blue. (A), (B), and (C) visualized in VMD. **(D, E, F)** Force on the cargo region over the course of each of the five trajectory replicates for the simulations without NBAR (D), with 10% surface coverage NBAR (E), and with 25% surface coverage. Dotted lines correspond to timepoints of rupture in representative trajectories from (A), (B), and (C).

By comparing frames from a representative trajectory for each surface coverage, we can observe how the geometry of the necks change as a function of the number of NBAR dimers, particularly at later time steps. Concurrently, we calculate the timeseries of the force exerted by the lipids on the cargo region for five replicates per surface coverage condition (Figure 6D). Each replicate is initialized with randomized placement of the NBAR dimers near the membrane surface. For example, at 175 10^6^CGts, the neck diameter decreases with increased surface coverage, as well as having a greater eccentricity (as shown in Figure S7). The comparative narrowness of the neck facilitates membrane hemifission in the plain membrane (Figure 6A, panel 4) at an average maximum force of 450 kcal/mol/nm (Figure 6D) before rupture. By contrast, the increased diameter of the neck at 10% and 25% NBAR dimer surface coverage (Figure S7A) lead to delayed hemifission (Figure 6B and 6C, panel 4) at larger average maximum forces of 490 kcal/mol/nm and 520 kcal/mol/nm, respectively (Figure 6E,6F).

Interestingly, in addition to allowing the membrane neck to sustain increased maximum forces before hemifission, we also note that the force plateaus post-rupture are also higher on average, indicating that the leaflet directly contacting the “cargo” surface is less strained after hemifission, and in the case of 25% surface NBAR coverage, held intact. This difference is attributable to the reorganization of lipids around the NBAR scaffold at the neck creating local lipid reservoirs that prevent premature rupture (Figure S8).

As noted previously, the NBAR dimers are enriched at the neck and excluded from the top half of the bud. This is due to their recruitment to the neck during bud formation, primarily (but not exclusively) from the membrane area associated with the bud (Figure S9A, S9B). There is a subsequent reorientation of the NBAR dimers associated with the neck antiparallel to the Z-axis (direction of bud motion) that is consistent with our previous observations on both straight and catenoid membrane tubes, generating the NBAR scaffold and associated lipid reservoir (Figure S9C, S9D). A full reorientation is not observed, likely due to the velocity of the cargo in the simulation being faster than the diffusive timescale of the NBAR dimers. However, it is interesting to note that the observed benefits of increased maximum force sustained after hemifission do not necessitate a full scaffold, as particularly exemplified at 25% surface coverage. These results are particularly notable for ultrafast endocytosis and fast-endophilin mediated endocytosis(2,53), two pathways that occur at timescales that preclude the total assembly of an endophilin scaffold. We contrast our hemifission results with the proposed friction driven scission (FDS) mechanism, where pore formation at the scaffold edge is the initiating event for scission(48). Rather, here we see a more continuous narrowing of the inner membrane until a hemifission intermediate is formed – often between scaffold layers. We note that in our CG simulations, the scaffold does not fix the underlying neck radius to be constant but rather can re-orient and respond to changes in tension and neck diameter, consistent with our other simulations reported earlier in this paper, thus emphasizing the dynamic interplay of membrane-mediated endophilin assembly formation.

## CONCLUSIONS

We have endeavored here to present a more complete picture for the remodeling capabilities of the NBAR-containing protein, endophilin. While the various mechanisms of endophilin-mediated membrane remodeling have been described as insertion (H0), scaffolding (BAR), and crowding (solvated unstructured and SH3 domains), up to this point they have not been directly compared altogether and on multiple membrane curvatures, particularly at the molecular (or quasi-molecular, i.e., CG) level. In particular, we emphasize the importance of insertion and scaffolding synergy in the complete NBAR unit, which is seen to be just as effective in remodeling as full length endophilin. Even at 10%, a relatively low surface coverage, we see the assembly of endophilin into string-like (flat membranes) or scaffold-like (membrane tubes) assemblies perpendicular to the membrane normal. We note that this low surface coverage is reasonably close to physiological expression levels and therefore more likely to be representative of cellular geometries.

Beyond the subdomain analysis, we have further explored the curvature sorting and generation ability to include systems with negative Gaussian curvature, such as catenoid tubes and Ω-shaped bud geometries. We have demonstrated at the molecular level the preference of endophilin for negative Gaussian membrane curvature. Our analysis reveals that NBAR assemblies locally sequester lipids to create “reservoirs” that help them to resist changes in membrane tension, and prevent hole formation during hemifission. Furthermore, these benefits arise even without a perfectly oriented scaffold, which may have implications for the recruitment and organization of downstream proteins such as dynamin(52,54,55).

While the hemifission method of membrane breaking does not agree with the previously-described friction-driven scission (FDS), we also note a few key differences such as the negative Gaussian curvature and rapid kinetics of scaffold assembly(48). In addition, we emphasize here the importance of modeling membranes with negative Gaussian as a difficult but crucial challenge for not only BAR domain proteins, but likely all curvature sensitive peripheral membrane proteins.

The CG simulations presented in this work provide insight on the possible behavior of NBAR-containing proteins such as endophilin and generate new hypotheses for future exploration and confirmation via experimental studies.

## AUTHOR CONTRIBUTIONS

JRB, PB, and GAV designed the initial project. JRB performed simulations and analyzed results with feedback from F-CT, GAV, and PB. JRB wrote the original draft with inputs and revisions from F-CT, PB, and GAV.

Conceptualization: JRB, PB, and GAV

Methodology: JRB

Investigation: JRB

Visualization: JRB

Supervision: PB, GAV, F-CT

Writing—original draft: JRB

Writing—review & editing: JRB, F-CT, PB, GAV

Funding acquisition: FCT, GAV, PB

## ACKNOWLEDGMENTS

JRB and GAV acknowledge funding from the National Institute of General Medical Sciences (NIGMS) of the National Institutes of Health (NIH) through NIGMS grant R35GM158238 (to GAV), the Chateaubriand Science and Technology Fellowship, and the University of Chicago France and Chicago Collaborating in The Sciences. F.-C.T. and P.B. are members of the CNRS consortium Approches Quantitatives du Vivant (AQV), Labex Cell(n)Scale (<u>ANR-11-889 LABX0038</u>) and Paris Sciences et Lettres (<u>ANR-10-IDEX-0001-02</u>). PB team is supported by the Fondation pour la Recherche Médicale (FRM) (FRM EQU202003010307) and by the European Union (ERC, PushingCell, #101071793). Views and opinions expressed are however those of the authors only and do not necessarily reflect those of the European Union or the European Research Council. Neither the European Union nor the granting authority can be held responsible for them. Simulations were performed using computing resources provided by the University of Chicago Research Computing Center (RCC), the Department of Defense High Performance Computing Cluster (HPCMP), and the National Science Foundation ACCESS cluster. JRB acknowledges Dr. Siyoung Kim and Dr. Zachary Jarin for thoughtful discussion and technical assistance.

