## Supplementary Materials for "Subdomains of Endophilin-NBAR Can Synergistically Drive Membrane Remodeling and Facilitate Controlled Membrane Scission"

**This PDF file includes:**

Table S1
Figs. S1 to S8

| System | Initial Membrane Size | # of Proteins | Production Time ($10^6$ CGts) |
|---|---|---|---|
| Flat Bilayer | | | |
| No Protein | 100 x 100 $nm^2$ | 0 | 100 |
| H0 | 100 x 100 $nm^2$ | 64 | 100 |
| BAR | 100 x 100 $nm^2$ | 32 | 100 |
| NBAR | 100 x 100 $nm^2$ | 32 | 100 |
| FL-endo | 100 x 100 $nm^2$ | 32 | 100 |
| Straight Tube | | | |
| No Protein | 30nm diam, 60nm len | 0 | 50 |
| H0 | 30nm diam, 60nm len | 48 | 50 |
| BAR | 30nm diam, 60nm len | 24 | 50 |
| NBAR | 30nm diam, 60nm len | 24 | 50 |
| Catenoid Tube | | | |
| No Protein | 30-60nm diam, 60nm len | 0 | 50 |
| NBAR | 30-60nm diam, 60nm len | 78 | 50 |
| Bud Generation | | | |
| No Protein | 150 x 150 $nm^2$ | 0 | 210 |
| NBAR (10%) | 150 x 150 $nm^2$ | 72 | 210 |
| NBAR (25%) | 150 x 150 $nm^2$ | 178 | 210 |

**Table S1.** Summary of CG simulations conducted in this study.

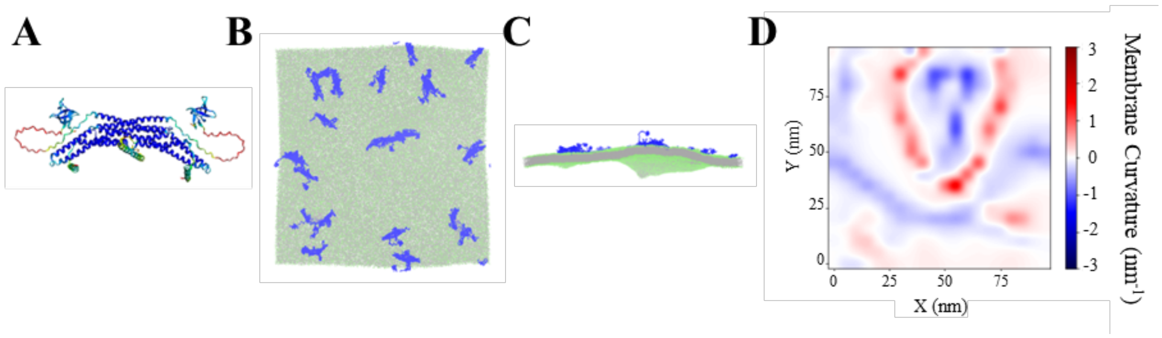


**Fig. S1.** Structure and remodeling of full length endophilin. **(A)** Structural representation of full length endophilin dimer, visualized with ChimeraX. **(B)** Top and **(C)** Side view snapshots of full length dimer CG systems after 100 $10^6$ CG timesteps (CGts) from random initial protein placement on $100x100nm^2$ membranes at 10% surface coverage. Dimers are represented in blue, with the lipid head sites shown in green and tail sites shown in silver, visualized with VMD. **(D)** Mean membrane curvature of upper (protein-bound) leaflet after 100 $10^6$ CGts. Membrane curvature calculated with MDAnalysis package MembraneCurvature.

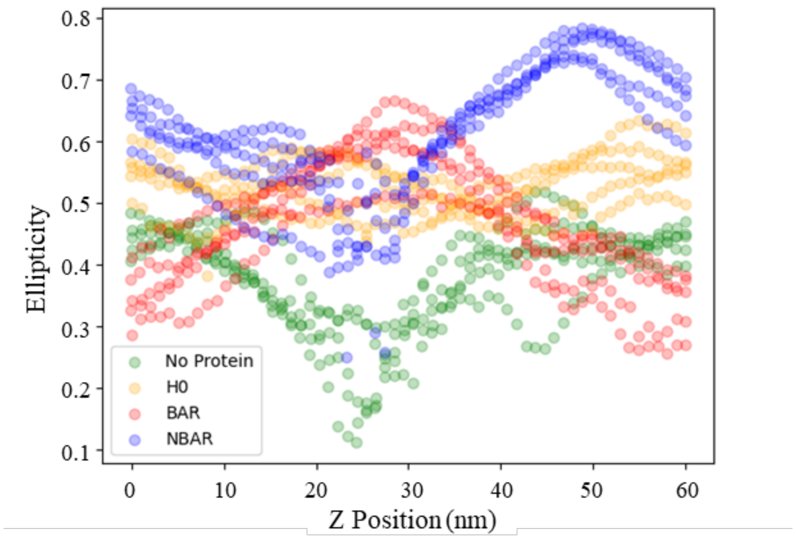


**Fig. S2.** Ellipticity, calculated as ratio of major and minor axes of initially straight tube slices averaged over every 1 nm in z-axis for membranes with no protein (green), with 24 NBAR dimers (corresponding to 10% surface coverage; blue), with 24 BAR dimers (red), and with 48 H0 helices (orange). Calculated for five evenly spaced time points from 45-50 $10^6$ CGts to show variability.

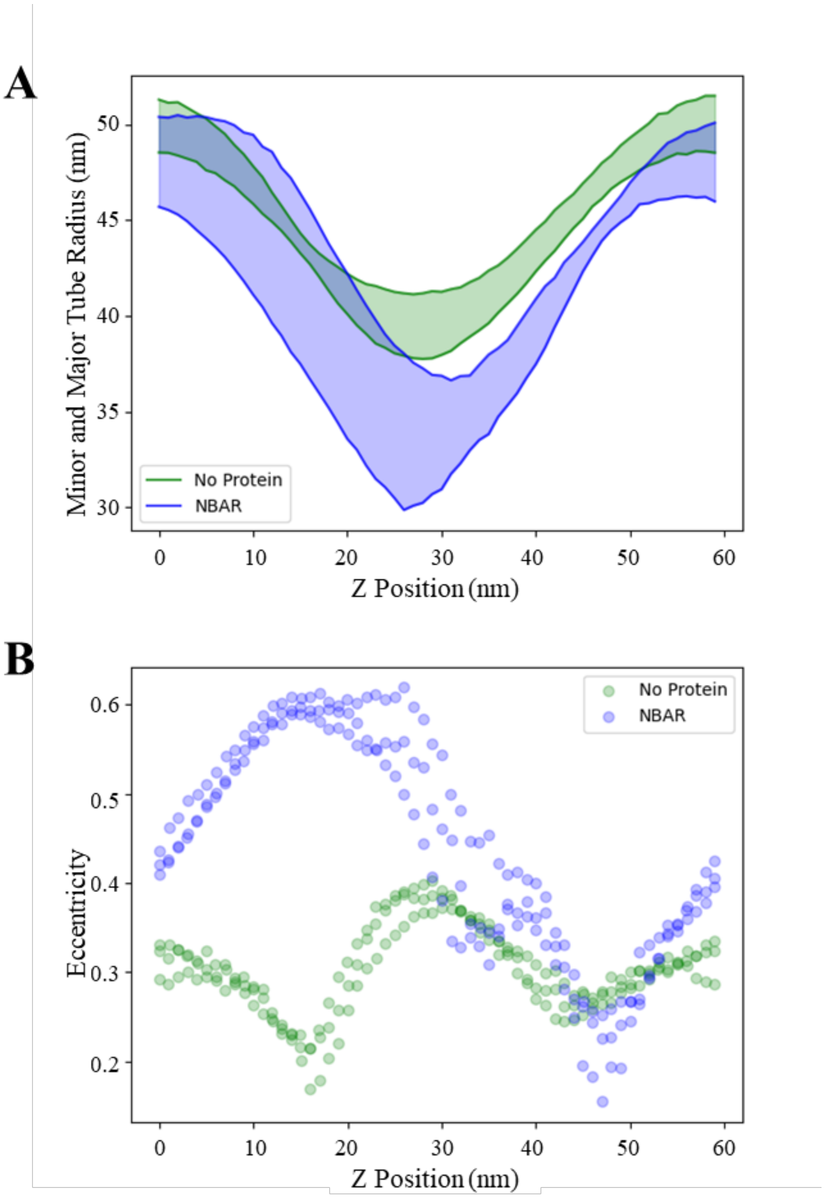


**Fig. S3.** Geometry changes in catenoid tubes after 50 $10^6$ CGts. (**A**) Minor and major tube radii in initially straight tubes after 50 $10^6$ CGts. Radii calculated based on CG tail bead positions subdivided into 1 nm wide slices for membranes with no protein (green) and with 24 NBAR dimers (corresponding to 10% surface coverage; blue). (**B**) Eccentricity as a function of z position based on ratio of minor and major tube radii for last 5 $10^6$ CGts in to show variance for membranes with no protein (green) and with 24 NBAR dimers (blue).

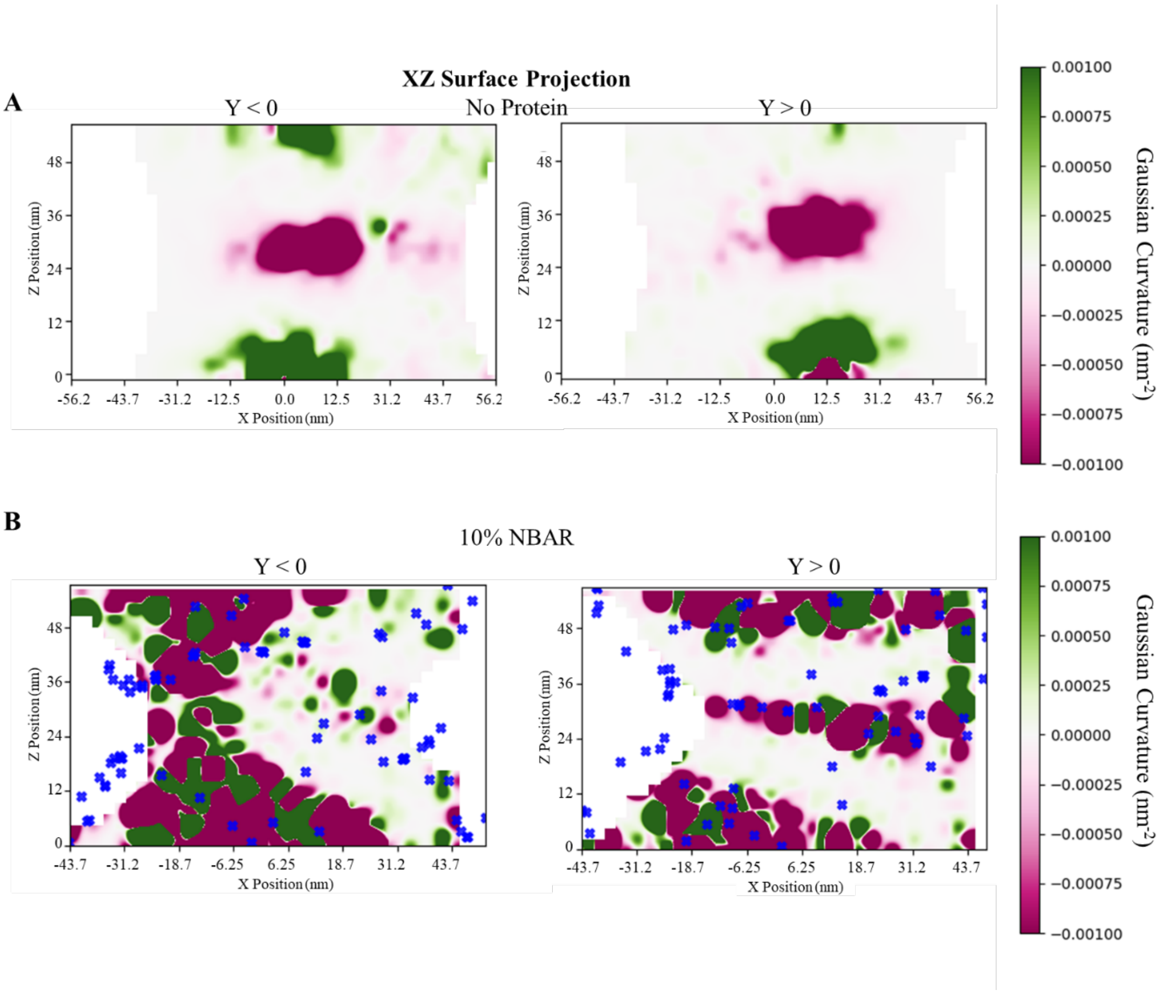


**Fig. S4.** Local Gaussian curvature projected onto xz plane after 50 $10^6$ CGts. **(A)** Back (Y < 0, *left*) and front (Y > 0, *right*) of catenoid tube membrane without NBAR protein, with Gaussian curvature calculated over 2.5x2.5nm$^2$ bins. **(B)** Back (Y < 0, *left*) and front (Y > 0, *right*) of catenoid tube membrane with 78 NBAR dimers (corresponding to 10% surface coverage), with Gaussian curvature calculated over 2.5x2.5nm$^2$ bins. NBAR dimer tip positions shown as blue x's. Gaussian curvature calculated with MDAnalysis package MembraneCurvature.

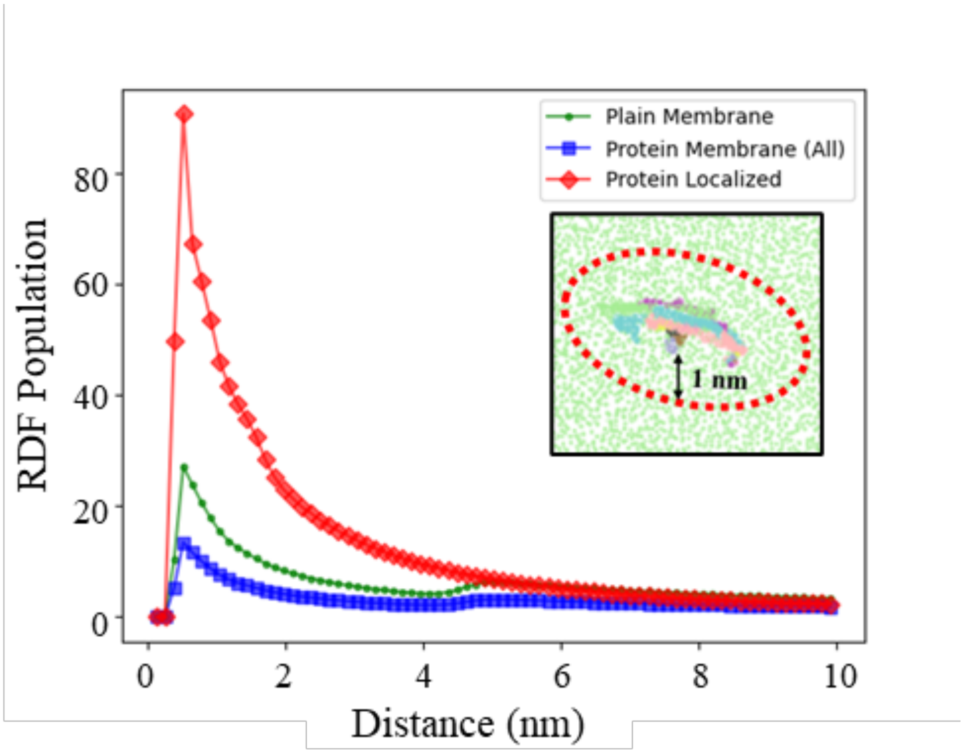


**Fig. S5.** Radial distribution function (RDF) in catenoid tubes, calculated from 45-50 $10^6$ CGts. RDF calculated using lipid head groups over entire membrane without protein (green dots), over entire membrane with 78 NBAR dimers (corresponding to 10% surface coverage; blue squares), or membrane area within local area (1 nm radius) of an NBAR dimer bead (red diamonds). *Inset*: visualization of 'protein localized' lipids within 1 nm radius of an NBAR dimer, generated with VMD.

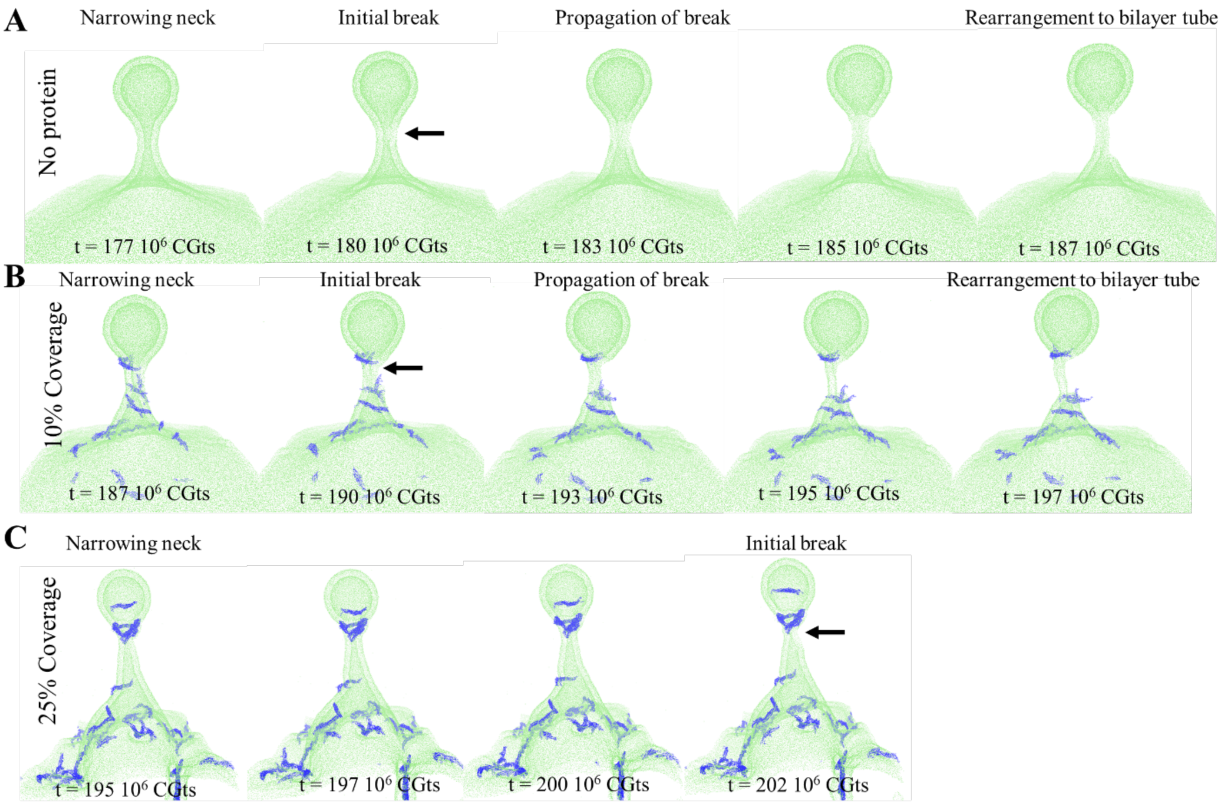


**Fig. S6.** Snapshots of transition from narrowing neck to hemifission intermediate from representative trajectories with **(A)** no protein, **(B)** 10% surface coverage of NBAR dimers, and **(C)** 25% surface coverage of NBAR dimers.

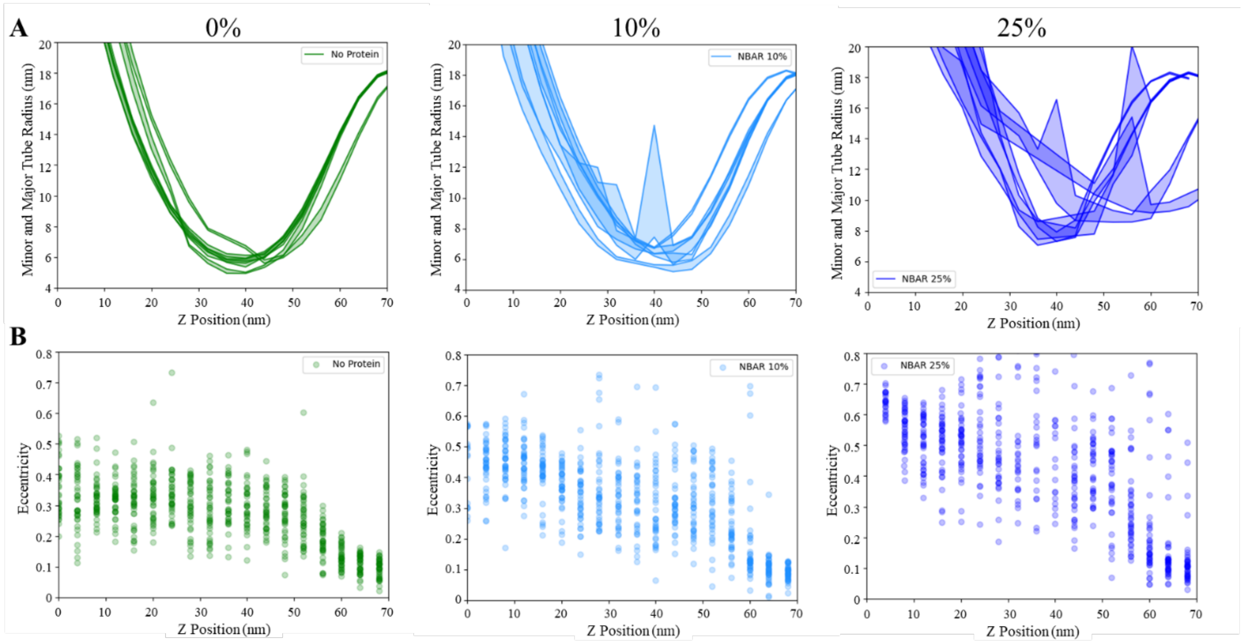


**Fig. S7.** Membrane geometry in bud formation, calculated from 170-175 $10^6$ CGts. (**A**) Minor and major tube radii in bud formation. Radii calculated based on CG tail bead positions subdivided into 4 nm wide slices for membranes with no protein (*left, green*), with 72 NBAR dimers (*center, light blue*), and with 178 NBAR dimers (*right, dark blue*). Fill is added between minor and major radii of each replicate for ease of visualization. (**B**) Eccentricity as a function of z position based on ratio of minor and major tube radii to show variance for membranes with no protein (*left, green*), with 72 NBAR dimers (corresponding to 10% surface coverage; *center, light blue*), or with 178 NBAR dimers (corresponding to 25% surface coverage; *right, dark blue*).

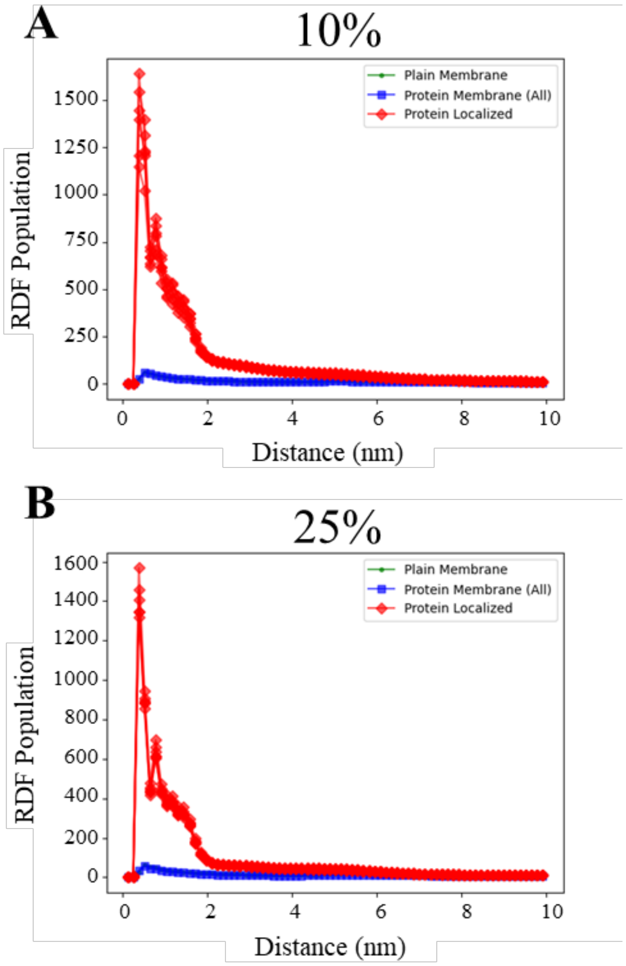


**Fig. S8.** Radial distribution function during bud formation, calculated from 170-175 $10^6$ CGts. **(A)** RDF calculated using lipid head groups over entire membrane without protein (green dots), over entire membrane with 72 NBAR dimers (corresponding to 10% surface coverage; blue squares), or membrane area within 1 nm of any NBAR dimer bead (red diamonds). Lines represent individual replicates. **(B)** RDF calculated using lipid head groups over entire membrane without protein (green dots), over entire membrane with 178 NBAR dimers (corresponding to 25% surface coverage; blue squares), or membrane area within local area (1 nm radius) of an NBAR dimer bead (red diamonds). Each line represents an individual replicate.

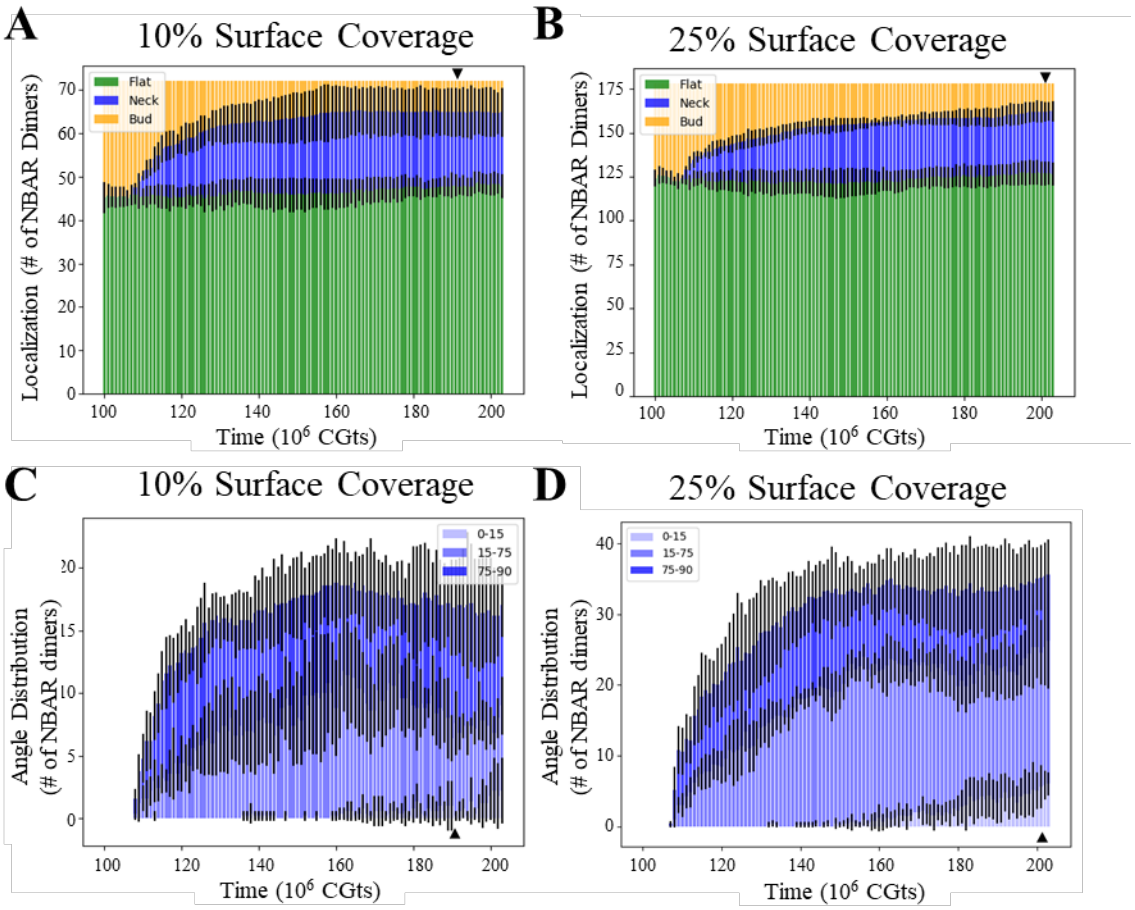


**Fig. S9.** Protein organization during bud formation. **(A, B)** Sorting of NBAR dimer center of masses based on Z-coordinate into original flat sheet, catenoid neck, and positive curvature bud regions for 10% (*A*) and 25% (*B*) surface coverage. Error bars represent standard deviation over all five replicates at same time point. **(C,D)** Distribution of NBAR dimer angle with Z-axis for dimers whose center of mass is sorted into catenoid neck region for 10% (*C*) and 25% (*D*) surface coverage. Arrowheads designate timepoint of rupture in representative trajectories in Figure 5. Error bars represent standard deviation over all five replicates at same time point.
